# The influence of body side and sex on neck muscle responses to left-frontal-oblique impacts

**DOI:** 10.1101/2020.12.04.406421

**Authors:** Andreas Mühlbeier, Kim Joris Boström, Marc H. E. de Lussanet, Wolfram Kalthoff, Cassandra Kraaijenbrink, Lena Hagenfeld, Jens Natrup, William H. M. Castro, Heiko Wagner

## Abstract

Low-velocity motor vehicle crashes frequently induce chronic neck disorders also referred to as whiplash-associated disorders (WAD). The etiology of WAD is still not fully understood. Women are affected more often and more severely than men. A frontal-oblique collision direction leads to WAD relatively frequently, but is poorly investigated as compared to rear-end or frontal collision directions. An oblique impact direction is assumed to strain the contralateral neck side more than the ipsilateral side, provoking an asymmetrical response pattern of the left and right cervical muscles. In this study, we examined the muscle reflex responses of the sternocleidomastoid, the paraspinal, and the trapezius muscles as well as the kinematic responses of 60 subjects during left-frontal-oblique collisions. Neither the reflex delay nor the peak head acceleration revealed significant sex differences. Thus, the elsewhere reported higher injury risk of females as compared to males cannot be explained by the electromyographic and kinematic results of this study. In females as well as in males the right muscle responses revealed shorter onset times than the left muscle responses. Thus, cervical muscles seem to be activated depending on the specific direction of the impact. Moreover, the difference between right and left muscle responses cannot be explained by a startle reflex. The movement of the head in relation to the torso (female mean ± SD: 92 ± 16 ms; male: 87 ± 16 ms) started almost simultaneously to the onset of the first muscle activation (right paraspinal muscles: female mean: 95 ms and 95%-CI 82 − 109; male mean: 95 ms and 95%-CI 84 − 108) indicating that the initial muscle activity seems not to be triggered by a stretch reflex in the PARA muscles.

## 1 INTRODUCTION

Whiplash-associated disorders (WAD) are among the most frequent medical complaints following low-velocity motor vehicle crashes (Chappuis and Soltermann, 2006; Graf et al., 2009). The etiology of WAD is still not fully understood and may depend on various occupant and crash related factors such as the passenger’s age (Harder et al., 1998), seating position (Berglund et al., 2003; Jakobsson et al., 2000; Jonsson et al., 2013), and sex (Dolinis, 1997; Harder et al., 1998; Richter et al., 2000; Berglund et al., 2003; Jonsson et al., 2013), as well as the vehicle’s change in velocity due to the impact (Δ*v*) (Kumar et al., 2005), and the impact direction (Berglund et al., 2003).

Based on 1001 individual medical examinations performed at the OFI Orthopädisches Forschungsinstitut Münster, we found that most of the WAD cases occur subsequent to rear-end collisions (56%). Interestingly, frontal-oblique collisions (16.7%) induced WAD more often than purely frontal (7.4%) or lateral collisions (7.7%, non-published data, see Mühlbeier et al. (2018)). Hence, it appears relevant to investigate frontal-oblique impact directions, which typically occur during intersection accidents and sliding collisions. So far, aligned impact directions (rear-end or frontal) have been well investigated. Such impacts strain the left and right cervical muscles in a similar manner, so that whiplash-associated symptoms, such as neck pain, are likely to occur symmetrically on both neck sides. However, during oblique collisions, the movement is more complex adding an asymmetric lateral and rotational component in the sagittal axis to the anterior-posterior movement of the head. Thus, one side of the neck may be strained more heavily than the other. Indeed, Kumar et al. (Kumar et al., 2004a,b) anecdotally reported that WAD patients who had exhibited an oblique collision direction tend to describe a unilateral neck pain. While little is known about the laterality of left and right individual cervical muscle responses to asymmetrical perturbation directions, it is plausible to assume that cervical muscles respond side-specifically to asymmetrical perturbations with a higher activation of the contralateral muscles. Following right anterolateral (Kumar et al., 2004a) and left anterolateral (Kumar et al., 2004b) impact directions, the splenius capitis muscle showed an increased muscle activation in that side of the muscle that was located contralateral to the impact.

Many studies have found a higher WAD risk in females than in males (Berglund et al., 2003; Jonsson et al., 2013; Dolinis, 1997). Females were found to show a significantly lower likelihood of recovery than males (Harder et al., 1998) and more females had longer lasting complaints as compared to males (Richter et al., 2000). The reasons for these sex differences are still unclear. Neck muscles can potentially alter the head and neck kinematics, because these muscles are active during the time interval of the collision in which the injury is likely to occur (Brault et al., 2000; Siegmund et al., 2002; Siegmund, 2011). Thus, it is plausible to approach the sex differences by comparing the neck muscle activity of females and males during a low-velocity collision. Siegmund et al. (2003a) reported larger normalized muscle response amplitudes of males in the sternocleidomastoid (SCM, *p* < 0.0001) and in the paravertebral (PARA, *p* < 0.01) muscles as compared to females. Furthermore, some studies showed an earlier neck muscle activation in females than in males (Siegmund et al., 2003a; Brault et al., 2000).

When analyzing the neuromuscular mechanisms in whiplash-like perturbations the kinematic as well as the muscular responses, and especially their temporal sequence, have to be considered. Head kinematics differences between sex have previously been found in studies considering rear-end collisions. Females were found to sustain greater (van den Kroonenberg et al., 1998; Schick et al., 2008; Linder et al., 2008; Carlsson et al., 2011) and earlier (Hell et al., 1999; Schick et al., 2008; Linder et al., 2008; Carlsson et al., 2011) peak head accelerations than males at impact velocities ranging from 4 to 9.5 km/h Δ*v*. Schick et al. (2008) also reported higher and earlier T1 x-acceleration peaks (females: 7.6 g/87 ms, males: 7.2 g/106 ms) and higher and earlier thorax x-acceleration peaks (females: 8.5 g/93 ms, males: 7.7 g/109 ms) for females compared to males. Carlsson et al. (2012) reported that the rearward peak x-displacement of the head relative to the torso occurred earlier for females compared to males at a Δ*v* of 8 km/h. The sequence of different movement onsets (torso, head, head to torso) and muscle activity onsets during a frontal-oblique collision provides important information about the temporal process of the collision events. Different onset timings between females and males could explain the higher injury incidence of females in low-velocity collisions.

Most of the available frontal-oblique data stem from investigations using post mortem human subjects (Forman et al., 2013) or crash test dummies (Pintar et al., 2007), which lack neuromuscular behavior. Furthermore, the experiments that investigated WAD mostly included rather small numbers of subjects. Therefore, we conducted an extensive study with 60 voluntary subjects exposed to low-velocity left-frontal-oblique impacts, to gain further insights into the neuromuscular mechanisms underlying whiplash-like perturbations that may cause the sex differences in the prevalence of WAD. In a within-subject study design, we varied several impact parameters to investigate their effect on the neck muscle response amplitude and delay, i.e. impact velocity difference Δ*v* (3 / 6 km/h), seating position (driver / front seat passenger), and deliberate pre-tension of the musculature (tense / relaxed) (Mühlbeier et al., 2018). In a between-subject design, we analyzed possible sex differences in the kinematic responses as well as in the neck muscle responses.

Based on the above presented literature we hypothesize that the neck muscles that are located contralateral to the impact direction show shorter muscle reflex delays than the ipsilateral neck muscles (H1). Further, we hypothesize that females show shorter neck muscle reflex delays than males (H2). Lastly, we hypothesize that the movement onsets of the head relative to the car as well as relative to the torso are earlier for female than for male subjects (H3) and that the peak acceleration of the head in relation to the car as well as in relation to the torso are higher for female than for male subjects (H4).

## 2 MATERIALS AND METHODS

### Subjects

Sixty-five healthy subjects participated in the experiment (Table 1). The data of five of them had to be excluded due to technical problems. All subjects signed an informed consent form that had been approved by the local ethics committee of the Department of Psychology and Sports Science of the University of Münster. All subjects possessed a driving license and were paid a nominal amount for the participation in the study. None of them was pregnant or suffered from chronic or acute neck or back pain. The individuals depicted in the figures of this manuscript have signed a consent for the publication of these case details.

**Table 1.** Mean (± SD) of the subject characteristics.

|  | Female | Male | *p*-Value |
| --- | --- | --- | --- |
| N | 26 | 34 |  |
| Age (years) | 30 ± 12 | 34 ± 14 | = 0.34 |
| Body mass (kg) | 65 ± 11 | 79 ± 12 | < 0.001 |
| Body height (cm) | 172 ± 6 | 183 ± 7 | < 0.001 |

### Experimental setup and procedure

#### Setup of the crash vehicle and the towing vehicle

The following methods description is partially adopted from Mühlbeier et al. (2018), as both studies are part of a bigger research project.

A Smart Fortwo (curb weight: 513 kg; year of manufacture: 2000) was chosen as the crash vehicle by virtue of its low weight. This model has two seats, one for the driver and one for the front seat passenger. The mass of the crash vehicle was further reduced by removing the engine and the gearbox (Figure doi: https://doi.org/10.1371/journal.pone.0209753.g001). Moreover, the A-pillars, the windshield, and parts of the steering wheel were removed to improve motion capture. To enable motion capture of the participants, a stiff metal construction was added to the front of the crash vehicle. The crash vehicle was drawn by a truck with the help of a towing mechanism that was fixed laterally to the truck. A metal bumper was added at the site of impact to protect the wheel housing. A tape switch on the surface of the pendulum buffer produced a flash light on each impact. A 12 volt truck battery provided power for the motion capture cameras and the electromyography (EMG) equipment.

The truck was prepared with a pendulum to impact the bumper of the crash vehicle. A foam block was used to soften the impact of the pendulum and to generate a realistic impact duration of approximately 0.1 s. The truck driver controlled a trigger mechanism that simultaneously released the crash vehicle and the pendulum.

#### Procedure and protocol

The subjects were measured in pairs. After being prepared with EMG electrodes and retro-reflective markers, the subjects adjusted their seats individually and connected their seat belts. One of the experimenters read out the experimental condition and the instructions. The driver was instructed to steer straight while the car was drawn by the truck. The pendulum was either dropped from a low or high position to cause a velocity change (Δ*v*) of either 3 or 6 km/h. An incident data recorder measured the acceleration of the crash vehicle during the impact to control for the proper change in velocity Δ*v*. The impact force was adjusted for each per pair of subjects according to their weights, such that the Δ*v* was within the required margin. Over all trials and subjects the mean ± SD Δ*v* was 3.15 ± 0.11 km/h and 6.2 ± 0.18 km/h, respectively. In the ‘tense’ task condition the instruction was:

> “Sie sitzen im Auto und Sie sehen, dass von links ein Fahrzeug kommt, das Sie übersehen hat. Eine Kollision ist unvermeidbar. Deshalb nehmen Sie kurz vor dem Aufprall eine angespannte Körperhaltung ein.”

In English this would read as: “You are driving in a car and see another car approaching you from the left side, which has overlooked you. A collision is unavoidable. Consequently, you adopt a tense posture briefly before the crash.” In the ‘relaxed’ condition the instruction was as follows:

> “Sie sitzen im Auto. Plötzlich kommt es unerwartet zu einer Kollision von links. Da die Kollision Sie unverhofft trifft, ist ihre Körperhaltung entspannt.”

In English: “You are driving in a car. Suddenly a collision hits you from the left side. Since the collision occurs unexpectedly, your posture is relaxed.”

The drivers were also instructed to keep their hands on the steering wheel, whilst the front seat passengers put their hands in their lap. Before each run, the vehicle was mounted to the truck and the pendulum was lifted to its initial position. The EMG recordings were started first, followed by the interior and exterior motion capture system that began their measurement synchronously on pressing a shared trigger button. The trigger signal was also recorded with the EMG recordings to synchronize them with the kinematics. The truck accelerated the crash vehicle to a speed of approximately 30 km/h. After reaching a location marked on the track, the truck driver pressed the release button to release both the crash vehicle and the pendulum. The pendulum impacted the bumper on the frontal-oblique direction side of the crash vehicle causing a resulting change in velocity of 3 or 6 km/h and a slight change of the driving direction to the right. As soon as the vehicle had assumed a straight and stable course, the driver stopped the vehicle.

At least ten runs were recorded for each pair of participants (Table 2). The first two runs were familiarization trials in tense posture with a Δ*v* of 3 km/h (first) and 6 km/h (second). After the familiarization runs, four experimental conditions with two different instructions and two different velocity changes were recorded in randomized order. After that the occupants switched their seating position. Again, the four experimental conditions were carried out in a randomized order. In case of a technical failure the run was repeated at the end of the four experimental runs. Since data processing was largely performed at a later time, a run was also repeated if the experimenters only suspected a technical failure (e.g. an impact outside the recording range of the external motion capture system). All 60 subjects experienced eight experimental conditions from all combinations of three independent variables: seating position (driver / front seat passenger), muscle tension (tense / relaxed), and change in velocity Δ*v* (3 km/h / 6 km/h). Ten of the 60 subjects participated in an additional experiment without a safety belt. Thus, the test protocol of these subjects had a different sequence and was extended by eight unbelted experimental conditions.

**Table 2.** Test protocol. (N = 60)

| Run No. | Driver | Front seat passenger | $\Delta v$ (km/h) | Muscle tone |
| --- | --- | --- | --- | --- |
| 1 | subject A | subject B | 3 | tense |
| 2 | subject A | subject B | 6 | tense |
| 3-6 | subject A | subject B | 3 or 6 | tense or relaxed |
| 7-10 | subject B | subject A | 3 or 6 | tense or relaxed |

### Electromyography

Surface EMG of the sternocleidomastoid (SCM), trapezius (TRA), and cervical paraspinal (PARA) muscles were bilaterally recorded with different EMG systems for the driver (DeMeTec, GJB Datentechnik GmbH, Germany) and front seat passenger (Biovision, Germany); both preamplified 1.000 times. As the posterior neck muscles are arranged in multiple layers, the term *paraspinal muscles* (PARA) actually represents the total muscle activity recorded at these electrodes (Mang et al., 2014). After the skin had been shaved and cleaned with medical abrasive paste (OneStep, H+H Medizinprodukte GbR, Münster, Germany), disposable Ag/Ag-Cl electrodes (H93SG, Covidien, Neustadt, Germany) with a circular uptake area of 0.5 cm in diameter were placed with an inter-electrode distance of 2.5 cm. The electrode placement was done according to Hermens et al. (1999) and Falla et al. (2002). For the SCM, the electrodes were attached along the sternal portion of the muscle, with the electrode center at 1/3 of the distance between the sternal notch and the mastoid process (Falla et al., 2002) (Figure doi: https://doi.org/10.1371/journal.pone.0209753.g002). For the TRA, the electrodes were situated halfway between the 7th cervical vertebra (C7) and acromion while the PARA-electrodes were located laterally to the spinal column directly under the hairline. The common ground electrode was affixed at C7. Tape was used to secure the electrodes. Muscle activity was measured bipolar at a sampling rate of 2000Hz (Software: ToM Erfassung, GJB Datentechnik GmbH, Germany).

### Kinematics

Retro-reflective markers were attached to the head, breast, shoulders, upper arms and forearms. The shoulder and breast markers were attached to the skin with double-sided and medical tape while the remaining markers were tightly fixed with straps to secure that markers stay in place. Except for the shoulder markers, all subject markers were grouped in clusters of at least three markers so that the 3D orientation of the body segment could be computed in addition to the 3D position. Additionally, markers were placed inside and outside the crash vehicle as well as on the seats. The four high-speed motion capture cameras (upgraded Oqus 300, Qualisys, Sweden) that were mounted at the front side of the car recorded the 3D movements of retro-reflective markers at 250 frames per second (fps). The distance from the starting point to the location of the crash was approximately 85 meters. At the crash location a series of ten further motion capture cameras (Oqus 400, Qualisys, Sweden, 250 fps) recorded the 3D kinematics of the vehicle around the time of impact, for about 14 m along the driving direction. Three markers were recorded by both the interior and exterior motion capture system to align the coordinates of the interior and the exterior motion capture recordings. A shared trigger construction ensured that the capture of the interior and the exterior motion capture systems were synchronized. Using this combined motion capture recording allowed us to compute the earth-bound motions of the participants.

### Data Analysis

The processing of the kinematic and EMG data was performed using MatLab (version 2017b, MathWorks, USA).

#### EMG data

In ten of the subjects, the activity of the tape switch caused an artifact that added a clean block pulse to the EMG signal of all six electrodes of each subject. In 14 further subjects, the tape switch added an artifact to only one specific electrode (SCM left). In all of these cases (10 subjects * 8 runs * 6 electrodes + 14 subjects * 8 runs * 1 electrode = 592 EMG signals; 21% of all EMG signals) the artifact was subtracted from the data (see Supplemental Material). In another 15 subjects, the tape switch caused an uneven artifact to EMG signals of varying electrodes. In these cases it was not possible to correct the EMG signals and the affected EMG signals were excluded from the statistical analysis (109 EMG signals, 4% of all EMG signals).

The 12V battery current was transformed to approximately 50Hz 220V by an on-board current transformer. The electromagnetic transformation sometimes caused a noise of approximately 50Hz plus harmonics that contaminated the EMG signals. To remove this noise, the EMG signals were filtered by a 50Hz periodic moving average that was subtracted from each signal individually (Levkov et al., 2005) (see Supplemental Material). A Fast Fourier Transformation (FFT) on the thus cleaned data showed a resulting smooth spectrum in the noise frequencies, even for the most severely perturbed signals (see Supplemental Material).

Consecutively, all EMG signals were high-pass filtered (4th-order Butterworth filter, 20Hz (De Luca et al., 2010)), rectified, and smoothed by a ± 10 samples moving average. The reflex onset was defined as the first sample where the signal exceeded 5 standard deviations above baseline activity, which in turn was calculated as the mean over a period of 300 ms before impact.

#### Kinematic data

The 3D positions and orientations of markers and rigid bodies (calculated using the clusters) were computed using the standard Qualisys analysis software (QTM version 2.16) and exported to Matlab files. The driving across the track caused slight deformations to the chassis. Consequently, the markers inside the cabin and on the seats were not entirely stationary, but made small coherent movements in the coordinate system of the cameras. The amplitude of these movements were in the order of a few mm. By subtracting the average movements of the cabin markers from the markers which were mounted to the subjects, it was possible to almost completely eliminate these movement artifacts from the subjects’ kinematics.

In a number of subjects, the movements that resulted from the crash caused a few markers to be obscured. Due to this, marker clusters of which the third marker disappeared temporarily were lost for the same period as well. This caused a fundamental problem, because such gaps in the position and orientation (even small ones) cannot be resolved by standard gap-filling. We devised a gap filling procedure as follows. A marker cluster has 6 degrees of freedom (DOF). Therefore, with one marker lacking, just a single parameter would have to be interpolated. Given cluster *ABX*, of which marker *X* disappears for some time, whereas vector 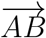 remains defined during continuously. Preceding and following the gap, the vector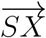, which is perpendicular to 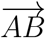 and the norm vector 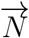 to *ABX* are defined. The 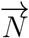 is then interpolated for the duration of the gap, using a 4th order polynomial interpolation and tails of 5 samples (using the Matlab `polyfit` function). The gap in *X* is then calculated as 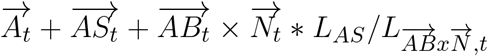, with subscript *t* for the time period of the gap in *X* and *L* being the vector length (i.e., the position of point *S* throughout the gap period, plus the cross product of vectors 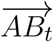 and the interpolated norm vector, scaled to the length of vector 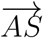. By this, effectively just a single parameter is interpolated (i.e., the orientation of plane ABX with respect to 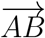) rather than all six parameters of the cluster. Tests on artificial gaps in trials when all markers remained visible throughout, showed that the error remained within the range of the measurement accuracy of the tracking system, even for artificial gaps across the 500 ms head movements following the impact. Gaps of up to 100 ms involving the shoulder markers were filled using the Matlab `polyfit` function, with a 4th order polynomial and a fit window with tails of 5 samples. The kinematic data was filtered using a 2nd order Savitzky-Golay filter with a window of 25 frames.

The movement onsets of 1) the torso (relative to the crash vehicle), 2) the head (relative to the crash vehicle), and 3) the head relative to the torso, respectively, were calculated from the corresponding rigid bodies (i.e., the average position of the marker cluster). The movement onset was defined as the moment when the tangential velocity exceeded a threshold of 10% of its maximum in the interval 300 ms following the impact. The peak velocity and peak acceleration of the head were defined as the maximum value, either in relation to the crash vehicle or in relation to the torso. As the impact direction in this investigation was frontal-oblique, we calculated the horizontal (x- and y-direction)torso and head kinematics in the direction corresponding to 45 degrees to the driving direction, perpendicular to the impact direction.

#### Reconstruction of the impact time

The slight deformation at the impact led to a small but highly specific and instantaneous motion of the tracking markers. The beginning of the impact was determined from the onset of this motion pattern with a tolerance of less than two frames (8 ms). To achieve this accuracy, each impact was determined at least two times independently; in cases when two different time samples had been determined, a third independent rating was made. Furthermore, the rating quality was checked by taking random trials from each measurement day for a further independent determination of the impact time. This manual determination method was far more accurate than various automatic procedures, such as a tape switch, and was the only method that was independent of the DeltaV.

#### Statistical Analysis

As some trials had to be excluded due to technical issues during the experiment, the final data structure became unbalanced, and so an N-way repeated-measures ANOVA was not viable. Instead, we performed a linear model (LM) analysis, and additionally, since the reflex delay was strictly positive, we calculated a generalized linear model (GLM) analysis. The latter were fitted with a logarithmic link function and gamma-distributed errors, while the LMs were fitted, as usual, with an identity link function and a Gaussian error distribution. The GLM fit converged better than the LM fit, in terms of both homoscedasticity and normality of the residual distribution as assessed by inspecting the diagnostic plots, so we will report the results from the GLM fit only. A certain extent of underdispersion could not be removed, but since underdispersion rather leads to a more conservative estimation of confidence intervals, we considered the fits sufficiently good for the purpose of this study.

The categorical factor levels have been symmetrically encoded (”effect coding”) with the weights ±1. The (G)LM models were each fitted with the maximum-likelihood method using the statistical software R with the package ‘lmerTest’ and the functions `lmer` for LM and `glmer` for GLM. The goodness of each model fit was assessed by inspecting diagnostic plots generated with the help of the DHARMA package, in particular Q-Q and residual plots.

We modeled the reflex delay as a continuous variable dependent on the following categorical variables: muscle type (`Muscle` = PARA, SCM, or TRA), muscle side (`Side` = left or right), Sex (`Sex` = male or female), change in velocity (`DeltaV` = 3 or 6 km/h), seating position (`Seat` = driver or front seat passenger), deliberate pre-tension of the musculature (`Tension` = tense or relaxed). We considered the unique subject ID as a categorical variable (`id` = 1,2,…, 60) that enters the model as random effect and nesting factor for within-subject variables. Because the EMG data were obtained using two different measurement systems with different output levels, we included an additional variable `EMGSys` to account for, and thus eliminate, these level differences.

The linear model was set up according to the formula

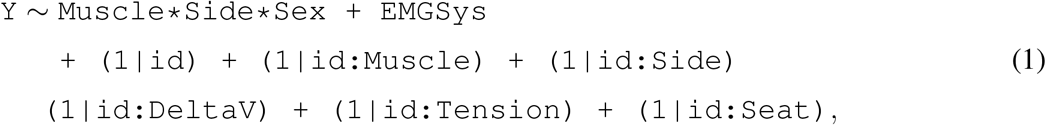

where `Y` stands for either the reflex delay or the reflex amplitude. Note that product terms of the form `A*B` are expanded to `1 + A + B + A:B`, where the constant term 1 represents the intercept, single terms `A` and `B` represent fixed main effects, colon-separated terms `A:B` represent interaction effects, and bracketed terms (`1|A`) represent random effects. In the presence of significant interactions between independent variables, the corresponding main effects may not have a straightforward interpretation and must be handled with caution.

In the present study, we focus on effects of the type of the measured muscles, the body side they are located, and the subject’s sex, so only `Muscle, Side` and `Sex` have been modeled as fixed effects, while `DeltaV, Tension`, and `Seat` were only modeled as random effects nested in the subject’s ID, hence generating within-subject variability. Because the variable `Sex` is a between-subject variable, there is no corresponding within-subject variability, hence no subject-nested random effect term.

To obtain p-values for the main and interaction effects, a type-3 ANOVA based on *χ*^2^-statistic was performed on each fitted model using the function `Anova` from the ‘car’ package. We chose to perform the more conservative type-3 ANOVA in view of potential interaction effects, and we based it on *χ*^2^-statistic rather than the more common F-statistic, because the former is more robust against potential non-normality of the data. The appropriate effect size parameter for *χ*^2^-tests is not 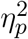 but Cramer’s *ϕ* (also known as Cramer’s *V*), which we calculated for each significant effect. The given interpretation of each effect size as *small*, *medium*, or *large*, depends on the degree of freedom (*df*) and was interpreted according to Cohen’s rule of thumb (Cohen, 1988) in the following way: For *df* = 1, a small/medium/large effect corresponds to values of *ϕ* around 0.1/0.3/0.5, and for *df* = 2, to values around 0.07/0.21/0.35.

Subsequent to each ANOVA, the functions `emmeans` and `pairs` from the package ‘emmeans’ were used to perform Tukey-adjusted post-hoc multiple-comparison tests.

## RESULTS

Exemplary EMG signals of the investigated cervical muscles during a left-frontal-oblique collision are given together with the acceleration profile in Figure 1. The acceleration of the crash vehicle occurred over approximately 100 ms (Figure 1, bottom). The mean SCM and TRA muscle onset was after this time window, but the perturbation has not necessarily to be over for the muscle to start its activity. The mean PARA muscle activity indeed begins already with the system still being under load.

**Figure 1.**
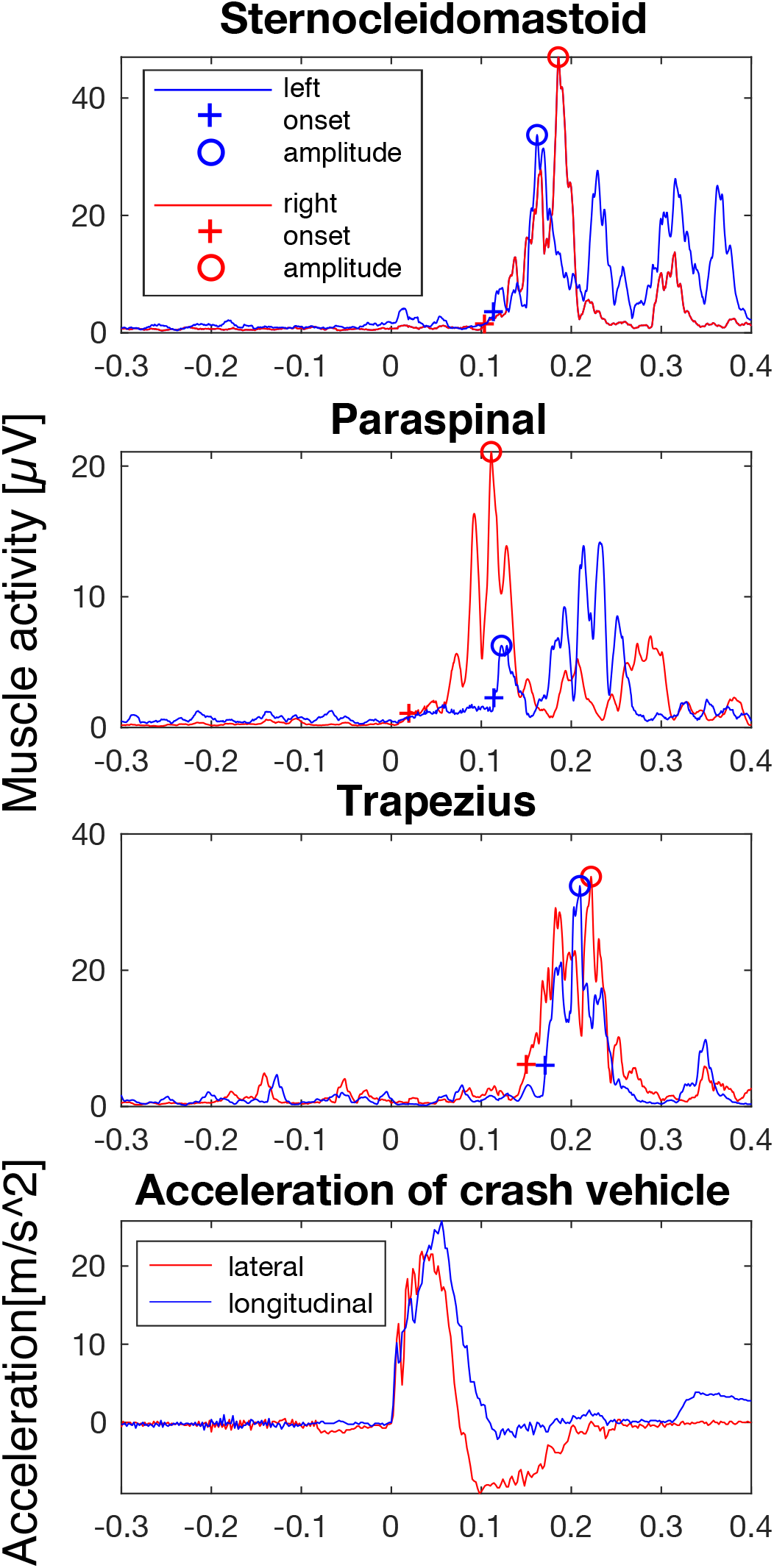
Exemplary EMG signals of an individual subject and acceleration of the crash vehicle. EMG signals of the left (blue) and right (red) sternocleidomastoid, paraspinal, and trapezius muscles of a male front seat passenger during an impact with a Δ*v* of 6 km/h and a relaxed pre-tension of the musculature. The muscle reflex onset and amplitude are indicated by a cross and circle, respectively. The time of 0 sec corresponds to the instant when the pendulum hit the bumper.

In line with H1, we found that in females as well as in males the right muscle responses revealed shorter onset times than the left muscle responses (Figure 2). No significant sex differences could be found for the reflex delay (*p* > 0.88, Table 3), so our hypothesis H2 is not supported by the data. Contrary to H3 we did not find an earlier female movement onset of the head in relation to the car whereas the movement onset of the head in relation to the torso of the female subjects was even significantly later compared to the males (Figure 3). No significant sex differences were found in the peak accelerations of the head, neither in relation to the car (*p* > 0.79, Figure 4, top right) nor in relation to the torso (*p* > 0.69, Figure 4, bottom right), hence H4 is not supported by the data.

**Figure 2.**
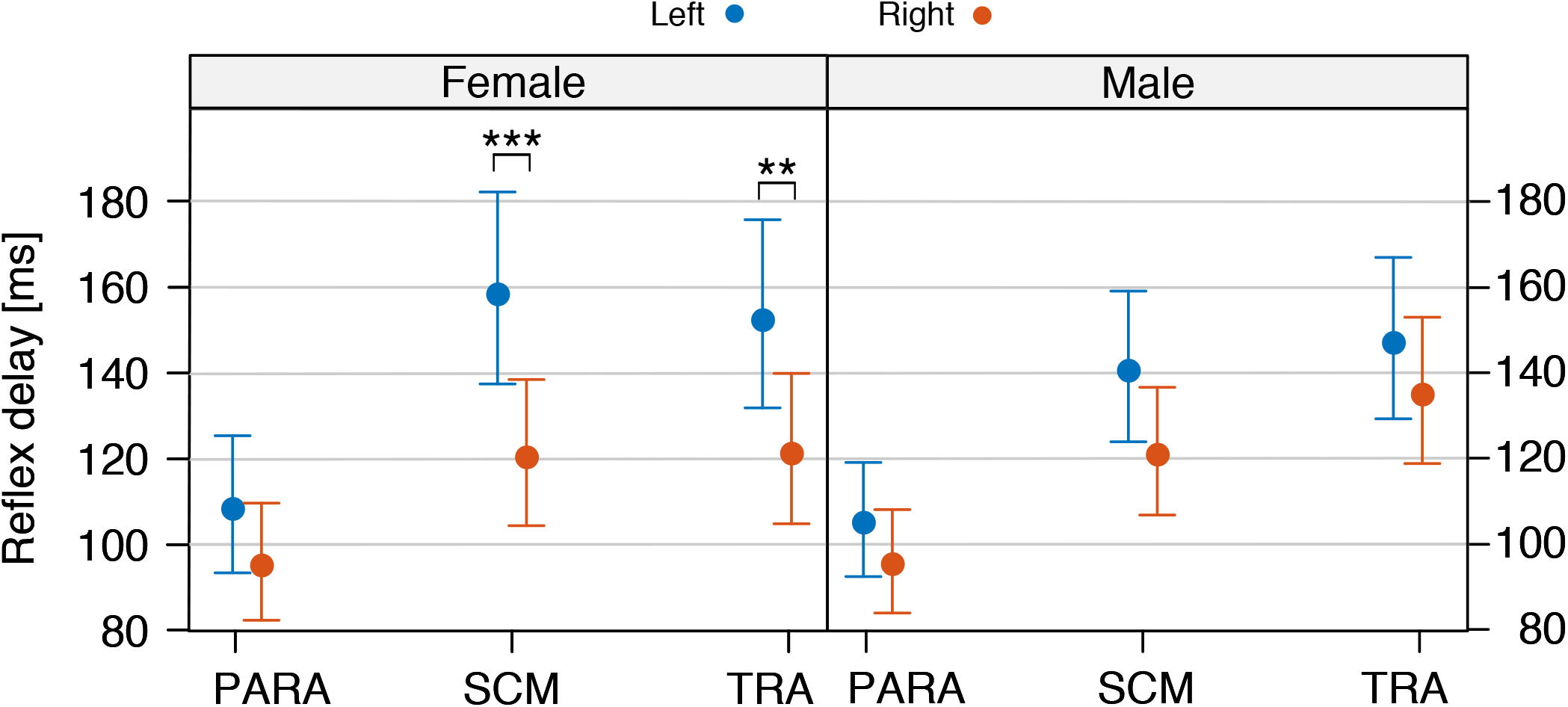
Reflex delay. Mean and 95%-confidence interval of the reflex delay [ms] of left (blue) and right (red) cervical paraspinal (PARA), sternocleidomastoid (SCM), and trapezius (TRA) muscles in 26 female (left) and 34 male (right) subjects.

**Table 3.**
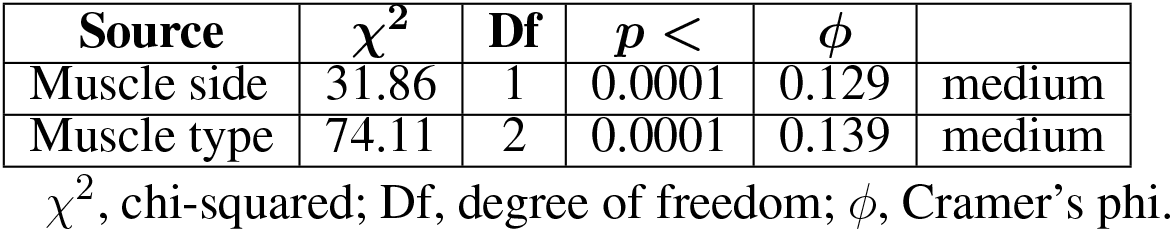
Main and interaction effects calculated by the statistical analysis of the reflex delay. Only significant effects are shown, p-values are rounded up to the next significant digit. Effect sizes are given as Cramer’s *ϕ* rounded to two significant digits, with a corresponding interpretation (small, medium, or large).

| Source | $\chi^2$ | Df | $p <$ | $\phi$ | |
| --- | --- | --- | --- | --- | --- |
| Muscle side | 31.86 | 1 | 0.0001 | 0.129 | medium |
| Muscle type | 74.11 | 2 | 0.0001 | 0.139 | medium |

$\chi^2$ , chi-squared; Df, degree of freedom; $\phi$ , Cramer's phi.

**Figure 3.**
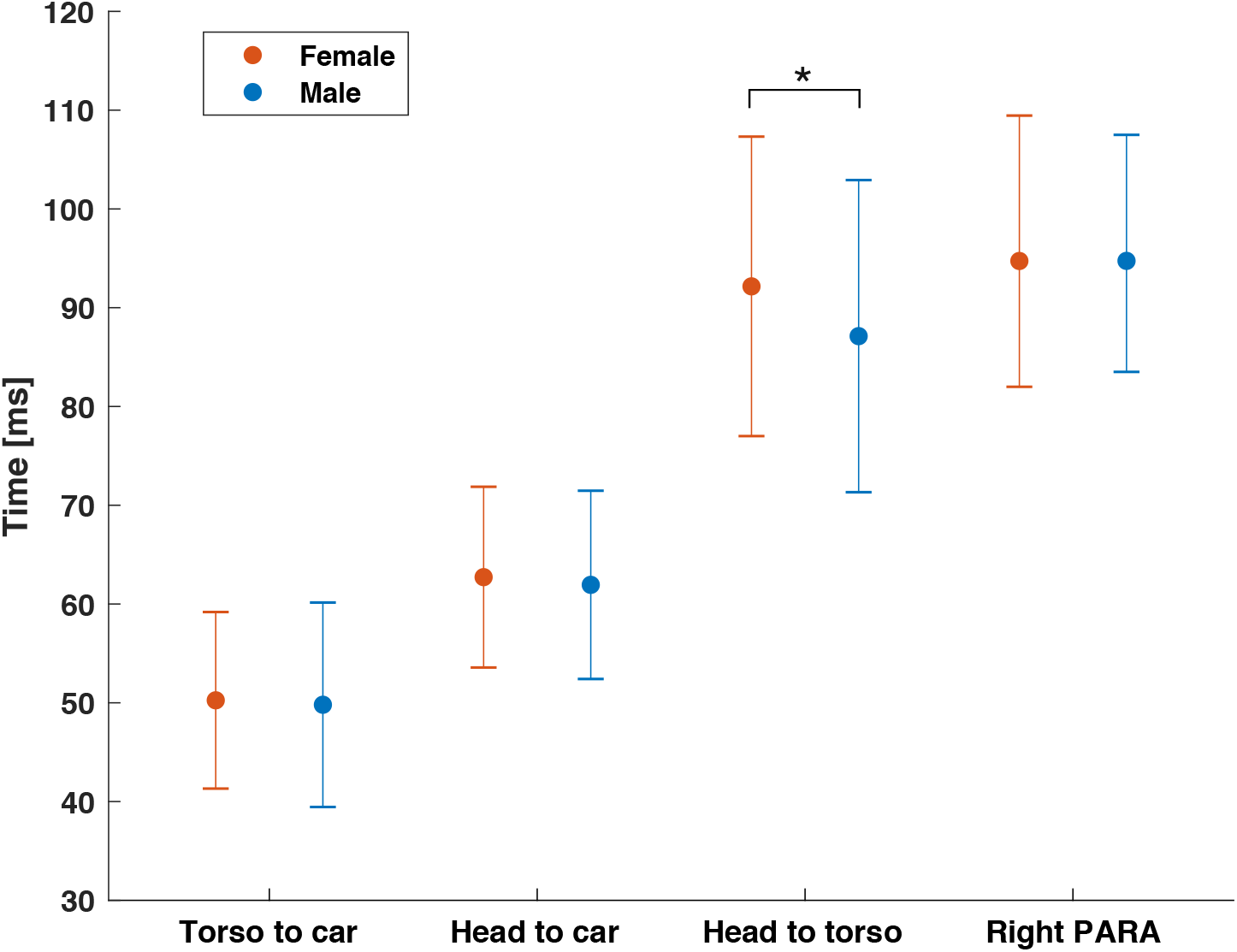
Kinematic and muscular onsets. Mean and SD of the movement onset of the torso in relation to the crash vehicle, the head in relation to the crash vehicle, the head in relation to the torso, and mean and 95%-confidence interval of the right cervical paraspinal muscles of 26 female and 34 male subjects.

**Figure 4.**
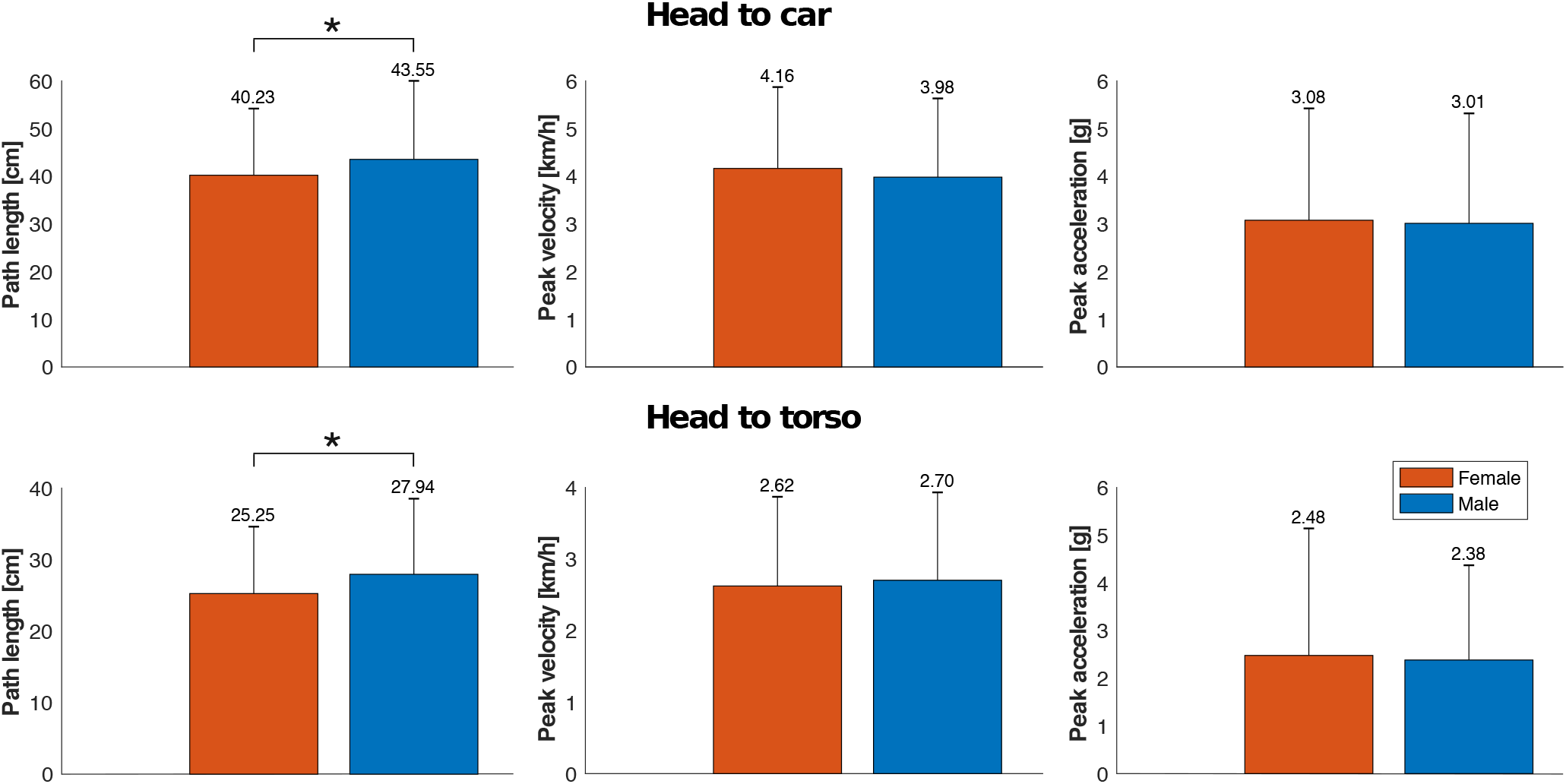
Path length, peak velocity, and peak acceleration of the head. Mean and SD of the path length (left), the maximum velocity (middle), and the maximum acceleration (right) of the head in relation to the crash vehicle (top) and in relation to the torso (bottom) of 26 female and 34 male subjects.

### Reflex delay

The statistical analysis (Table 3) revealed no significant main effect of the subject’s sex (*p* > 0.88) on the reflex delay (Figure 2). There was, however, a significant main effect of the muscle side (*p* < 0.0001) with shorter reflex delays on the right side of the body. The post-hoc tests showed significantly shorter reflex delays for the right SCM and TRA muscles of female subjects compared to their left counterpart.

### Movement onsets

The analysis showed an ascending sequence of movement onsets in the upper body. For female as well as for male subjects the torso started to move in space at 50 ms, about 12 ms before the head movement began (Figure 3). The movement onset of the head in relation to the torso at about 90 ms was followed by the first active neck muscles at about 95 ms. The statistical analysis revealed that the subject’s sex neither had a significant main effect on the movement onset of the torso in relation to the crash vehicle (*p* = 0.74), nor on the movement onset of the head in relation to the crash vehicle (*p* = 0.52). The movement onset of the head in relation to the torso was significantly earlier for male subjects compared to female subjects (*p* < 0.05).

### Head displacement, peak head velocity, and peak head acceleration

The GLM analyses revealed significant main effects of the sex on the path length of the movement of the head relative to the car (*p* < 0.05, Figure 4, top left) as well as to the torso (*p* < 0.05, Figure 4, bottom left) whereas there were no significant main effects of the sex on the maximum velocity nor on the maximum acceleration of neither the head relative to the car nor to the head relative to the torso.

## DISCUSSION

The purpose of this study was to examine the left and right neck muscle responses as well as the kinematic responses of females and males during low-velocity left-frontal-oblique collisions, to gain further insights into the neuromuscular mechanisms potentially related to whiplash-associated disorders (WAD).

In females as well as in males the PARA muscles were active first. This may be caused by the head’s primary movement towards the incoming pendulum from the left-frontal side as the dorsal PARA muscles are suited best to counteract this cervical flexion. The following rebound movement of the head towards the seat corresponds to an extension movement in the cervical joints and may be counteracted by the ventral SCM muscles which start their activity about 30–50 ms later than the PARA muscles.

The right muscle responses had shorter onset times as compared to the left body side. In investigations using aligned impact directions it has frequently been proposed that the neck muscle activity may be evoked by an (acoustic) startle reflex (Blouin et al., 2006; Mang et al., 2012, 2014, 2015). The startle reflex is known for its bilateral mechanism affecting both muscle sides equally and symmetrically (Yeomans et al., 2002). As we found differences between left and right neck muscle responses, it seems unlikely that the neck muscle reflex activity during low-velocity collisions is exclusively evoked by a startle reflex. Rather, the asymmetry of the left and right muscle responses is caused by the mechanical asymmetry of the left-frontal-oblique impact direction and is thus probably elicited by motion dependent reflexes such as a stretch reflex of neck muscles (Siegmund and Brault, 2000; Siegmund et al., 2008; Siegmund, 2011) or a vestibular reflex (Forssberg and Hirschfeld, 1994; Siegmund et al., 2008; Siegmund and Blouin, 2009; Siegmund, 2011). An additional indication for such an asymmetric muscular strain are the anecdotal reports of unilateral pain perception following an asymmetrical perturbation of the neck by the subjects participating in the studies of Kumar et al. (2004a,b). Still, the participation of the startle reflex as one of various neuromuscular mechanisms is possible as muscle responses may well result from a combination of several efferent signals of different origin.

Ito et al. (1997) investigated the earliest neck muscle responses of labyrinthine-defective (LD) subjects and healthy controls to a sudden fall of the head while laying supine on a horizontal surface. The healthy controls’ SCM muscles were active 24 ms after perturbation onset, whereas the LD subjects who could not rely on vestibular reflexes showed the first SCM muscle activation at 67 ms. The authors assumed that the initial EMG response in the LD subjects was a stretch reflex whereas the earlier EMG response in healthy subjects was a vestibular reflex. The relative movement between head and torso is the condition for the occurrence of a stretch reflex in the neck muscles. The onset of the head movement relative to the torso (females: 92 ms, males: 87 ms) was only slightly earlier than the onset of the right PARA muscles (females: 95 ms, males: 95 ms) which were active first in this investigation (Figure 3). Due to this short interval between head-torso movement and muscle response onset, a monosynaptic stretch reflex in the PARA muscles is unlikely to be the trigger of the initial muscle activity. However, it may still contribute to the muscle activity in later reflex components. The movement of the head in relation to the crash vehicle started about 30 ms earlier than the first muscle activity (Figure 3) providing enough processing time for a vestibular reflex to occur (Ito et al., 1997). Thus, the early neck muscle activity seems not to be initiated by the stimulation of the proprioceptors in the cervical spine due to the relative head-torso movement (cervical extension or flexion, lateral bending, etc.), but rather by the stimulation of the vestibular organs due to the earlier movement of the head in space.

A potential habituation of the muscle responses due to repeated exposures may lead to an attenuation of the muscle response amplitudes (Blouin et al., 2003; Siegmund et al., 2003b). However, a statistical comparison between the two familiarization trials (’Δ*v*=3 km/h tense’ and ’Δ*v*=6 km/h tense’) with the two subsequent experimental conditions of ’Δ*v*=3 km/h tense’ and ’Δ*v*=6 km/h tense’ revealed no significant effects on the reflex amplitude nor on the reflex delay. Possibly, the study design with a randomized sequence of experimental conditions that was used in this investigation prevented the CNS to adopt the muscular responses to a specific load.

While numerous studies have found sex-specific differences in WAD risk (Berglund et al., 2003; Jonsson et al., 2013; Dolinis, 1997), likelihood of recovery (Harder et al., 1998), or duration of complaints (Richter et al., 2000), only few investigations examined differences in the muscular responses of female and male subjects, probably because participant numbers were often not sufficient. Brault et al. (2000) and Siegmund et al. (2003a) investigated neck muscle responses of 40 and 66 subjects to rear-end collisions, respectively, and found earlier female than male cervical muscle onset times. In contrast to these results, our study revealed no significant sex-related differences for the reflex delay (*p* > 0.79). Anthropometric differences between male and female subjects have also been presumed to affect WAD (Vasavada et al., 2008). However, a separate statistical analysis, where subject height and weight were added as covariates, revealed that neither the height (*p* = 0.72) nor the weight (*p* = 0.78) had an influence on the reflex delay.

Contrary to the results of van den Kroonenberg et al. (1998); Schick et al. (2008); Linder et al. (2008); Carlsson et al. (2011) we did not find greater peak head accelerations or peak head velocities of female as compared to male subjects (Figure 4, right). There were small but significant sex differences in the path length of the head movement (Figure 4, left). However, the peak head acceleration is considered more appropriate to assess the genesis of WAD (Siegmund and Blouin, 2009). In contrast to Hell et al. (1999); Schick et al. (2008); Linder et al. (2008); Carlsson et al. (2011) who reported earlier kinematic responses of females, our analyses demonstrated not significantly different (torso to car and head to car, Figure 3) or even slightly earlier movement onsets of the males (head to torso: female mean ± SD: 92 ± 16 ms; male: 87 ± 16 ms, *p* > 0.05). Altogether, it is highly questionable if these small kinematic differences between female and male subjects can account for the reported sex differences in WAD risk.

## CONCLUSION

This study was the first to compare neck muscle and kinematic responses between males and females following frontal-oblique low-velocity impacts. As the neck muscle reflexes of female compared to male subjects are indistinguishable with respect to response delay, our data do not provide an explanation for the elsewhere reported higher injury risk of females as compared to males. Moreover, the results of this study provide a more precise picture of the neuromuscular mechanisms that may be involved in the genesis of WAD. Impacts from left-frontal-oblique directions evoke asymmetrical neck muscle responses with shorter reflex delays in the muscles opposing the side of the impact. Thus, the difference between muscle responses on the left and right side cannot be explained by a startle reflex alone. A monosynaptic stretch reflex of the cervical paraspinal muscles is unlikely as the initial neuromuscular mechanism triggering the muscle response as the head-torso movement that is required for the stretch of PARA muscles started only 3-8 ms before the onset of the right PARA muscles. Rather, the initial neck muscle activity may have been evoked by the stimulation of receptors in the inner ear (e.g. otoliths) due to the earlier movement of the head in space.

## ETHICS STATEMENT

This study was carried out in accordance with the recommendations of the ethics committee of the Institute of Psychology and Sports Science of the University of Münster (2015-44-HW) with written informed consent from all subjects. The protocol was approved by the the ethics committee of the Institute of Sports Science and Psychology of the University of Münster.

## AUTHOR CONTRIBUTIONS STATEMENT

All authors, especially WC and HW, made substantial contributions to the conception and the design of this study. WK, KB, AM, ML, HW, CK, JN and LH conducted and acquired the data of the experiment. KB, ML, and WK analyzed the kinematic data whereas AM evaluated the EMG data. KB performed the statistical analysis. AM wrote the first draft of the manuscript while all other authors made essential contributions by commenting and improving the final version. All authors approved the final version to be published, and are accountable for all aspects of the work.

## ACKNOWLEDGEMENTS

A special thanks goes to the entire team of the CTS (crashtest-service.com GmbH, Amelunxenstraße 30, 48167 Münster) for their competent technical support and the fruitful cooperation in the conducting of the experiment. Also we would like to thank Dr. Manfred Becke for his help in the planning of the experiments.

## SUPPLEMENTAL DATA

### DATA AVAILABILITY STATEMENT

The data supporting the conclusions of this manuscript will be made available by the authors to any qualified researcher.

