## Supplementary figures and images for "The influence of body side and sex on neck muscle responses to left-frontal-oblique impacts"

### S1_Fig

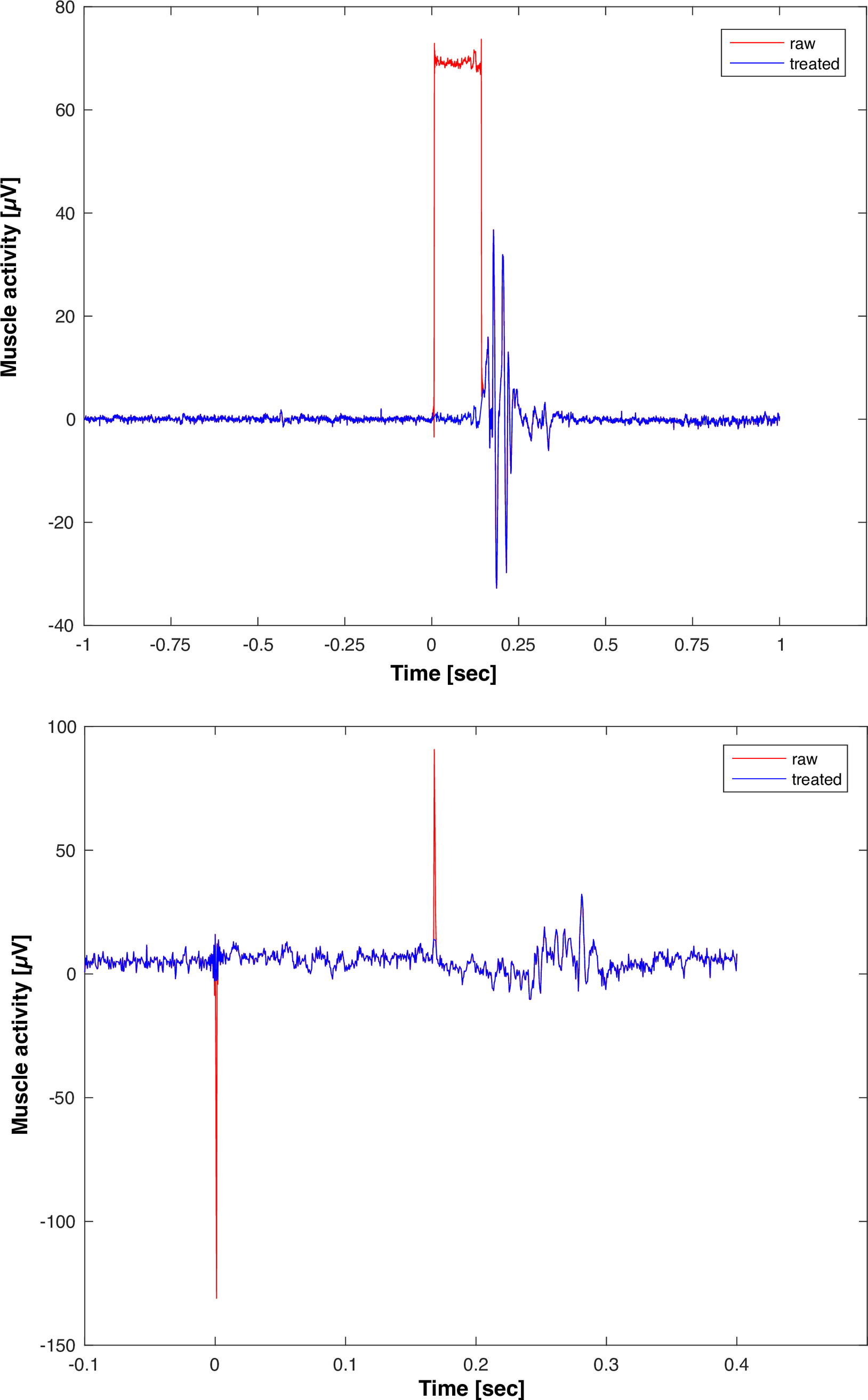

### S2_Fig

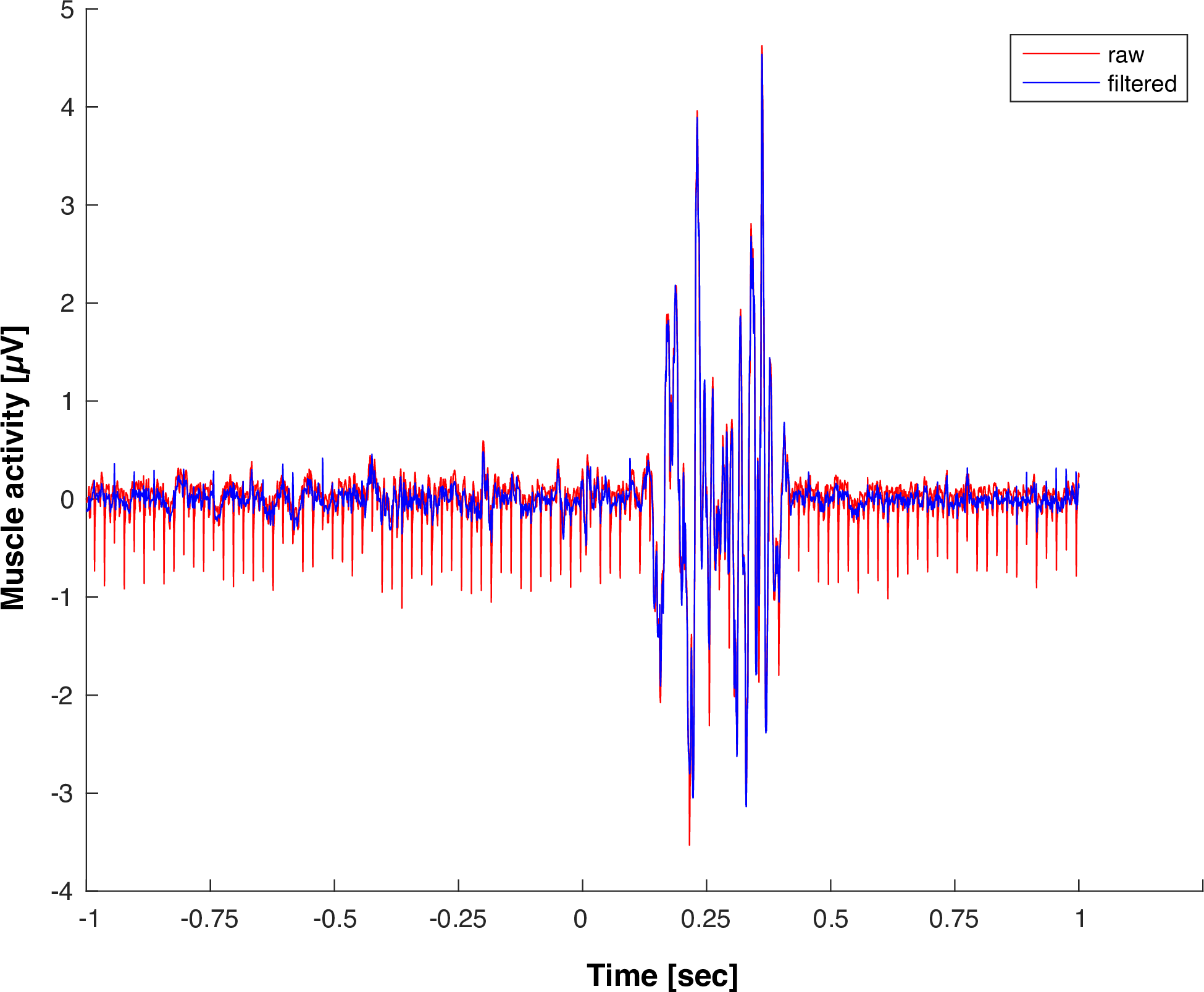

### S3_Fig

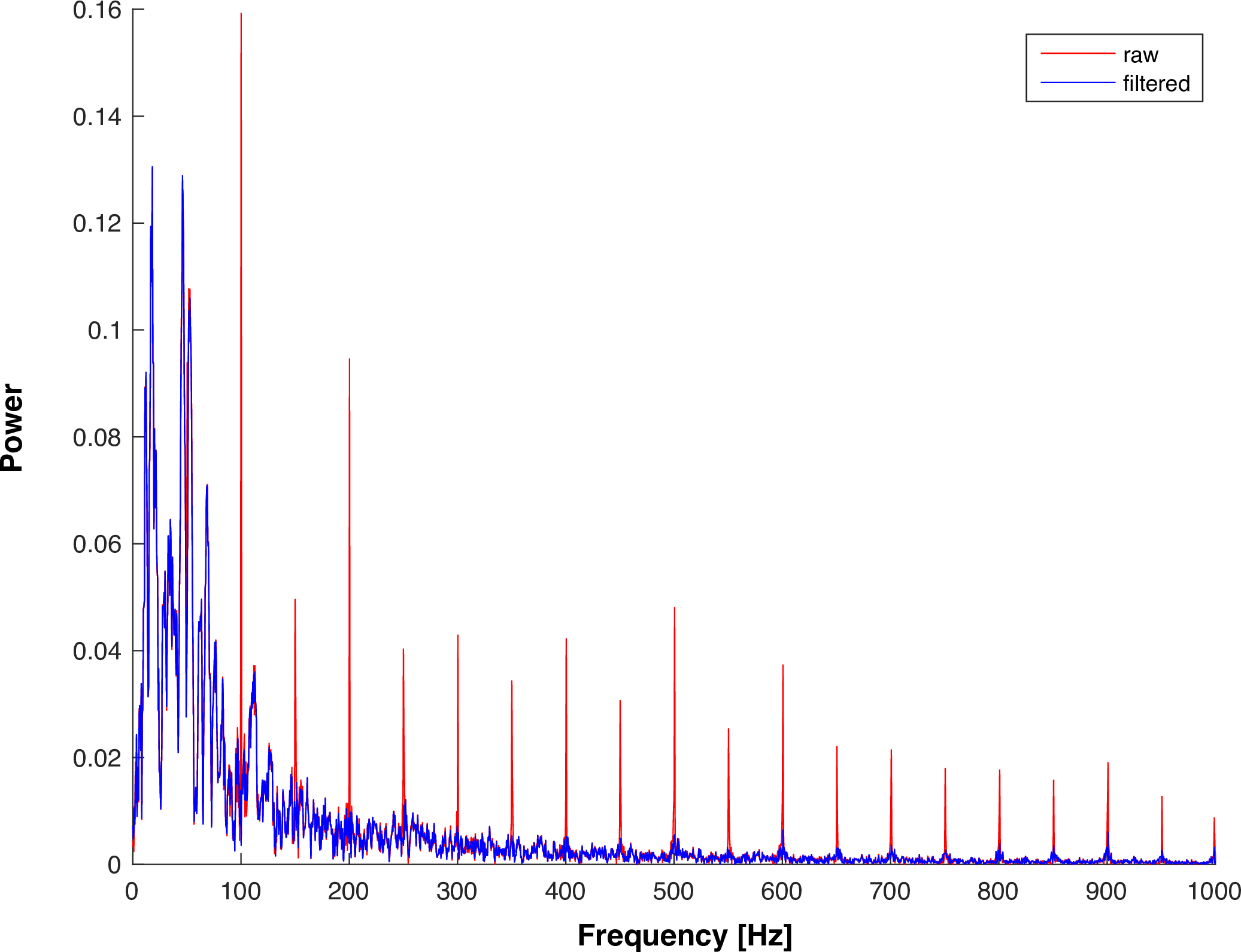

### S4_Fig

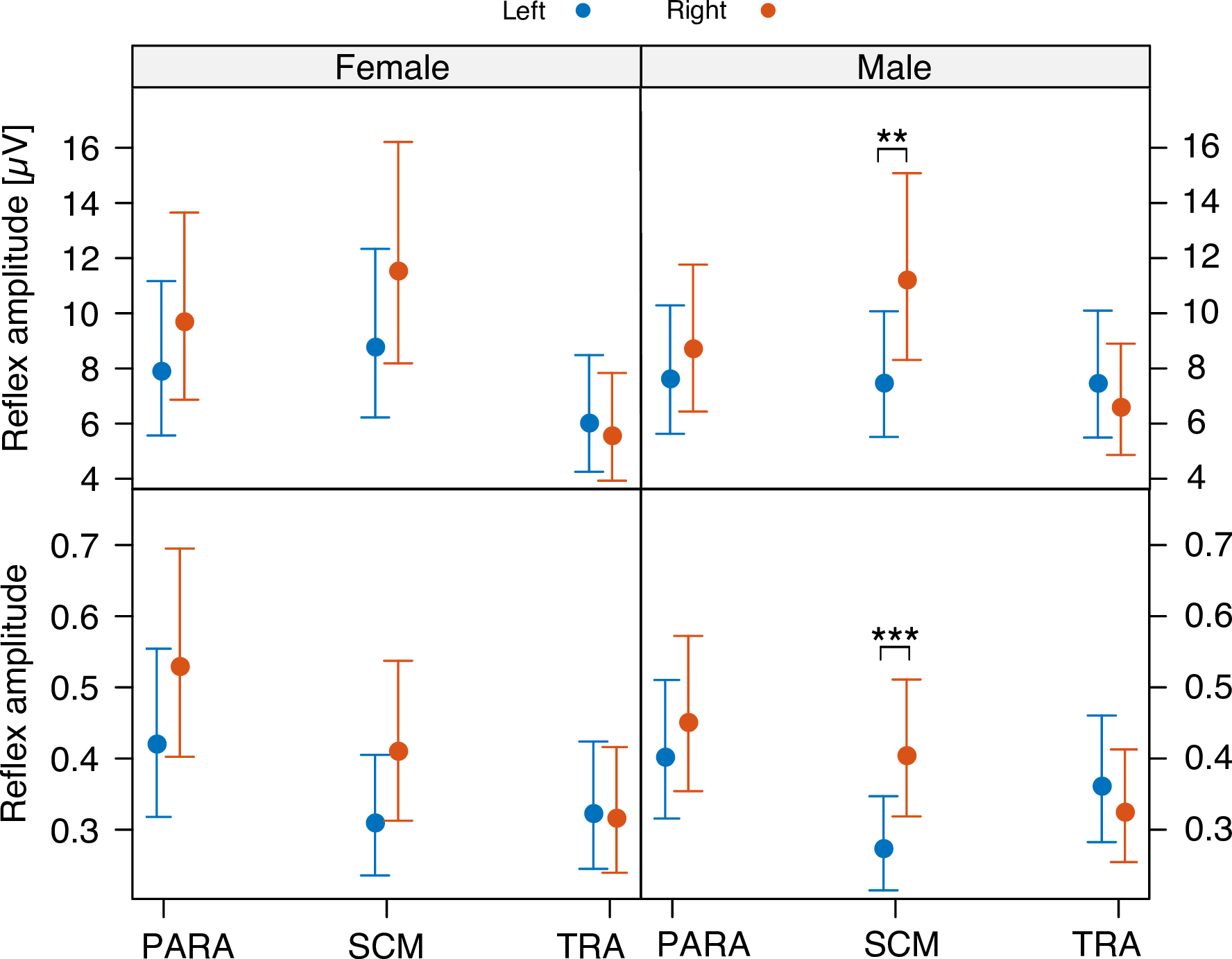
